# UPREGULATION OF SUPEROXIDE DISMUTASE 2 BY ASTROCYTES IN THE SIV/MACAQUE MODEL OF HIV-ASSOCIATED NEUROLOGIC DISEASE

**DOI:** 10.1101/2020.05.05.078691

**Authors:** Michelle N. Sullivan, Samuel A. Brill, Lisa M. Mangus, Yea Ji Jeong, Clarisse V. Solis, Audrey C. Knight, Carlo Colantuoni, Gizem Keceli, Nazareno Paolocci, Suzanne E. Queen, Joseph L. Mankowski

## Abstract

HIV-associated neurocognitive disorders (HAND) remain prevalent despite implementation of antiretroviral therapy (ART). Development of HAND is linked to mitochondrial dysfunction and oxidative stress in the brain; therefore, upregulation of antioxidant defenses is critical to curtail neuronal damage. Superoxide dismutase 2 (SOD2) is a mitochondrial antioxidant enzyme essential for maintaining cellular viability. We hypothesized that SOD2 was upregulated during retroviral infection. Using a simian immunodeficiency virus (SIV)-infected macaque model of HIV, quantitative PCR showed elevated *SOD2* mRNA in cortical gray (GM, 7.6-fold for SIV vs. uninfected) and white matter (WM, 77-fold for SIV vs. uninfected) during SIV infection. Further, SOD2 immunostaining was enhanced in GM and WM from SIV-infected animals. Double immunofluorescence labeling illustrated that SOD2 primarily co-localized with astrocyte marker glial fibrillary acidic protein (GFAP) in SIV-infected animals. Interestingly, in ART-treated SIV-infected animals, brain *SOD2* RNA levels were similar to uninfected animals. Additionally, using principal component analysis in a transcriptomic approach, *SOD2* and *GFAP* expression separated SIV-infected from uninfected brain tissue. Projection of these data into a HIV dataset revealed similar expression changes, thereby validating the clinical relevance. Together, our findings suggest that novel SOD2-enhancing therapies may delay the onset or reduce severity of HAND seen in ART-treated HIV-infected patients.

## Introduction

More than 37 million people in the world are infected with HIV (1), and nearly half of these individuals may develop HIV-associated neurocognitive disorders (HAND), including HIV-associated dementia (HAD), minor neurocognitive disorders (MND), and asymptomatic neurocognitive impairment (ANI) (2). Although the implementation of antiretroviral therapy (ART) has reduced mortality associated with HIV infection, HAND remains prevalent (2-4). Mitochondrial dysfunction and oxidative stress are known hallmarks of neurodegenerative diseases (5) and have been implicated as key factors in the development of peripheral neuropathy following HIV and simian immunodeficiency virus (SIV) infection (6-8). Therefore, investigation of mechanisms that preserve mitochondrial function and reduce oxidative damage is critical for development of therapies for HAND.

Mitochondria are the primary source of energy for cells in the brain (9). While other brain cells, like astrocytes, have alternative sources of ATP generation, neurons almost exclusively rely on mitochondrial oxidative phosphorylation (10). Byproducts of mitochondrial respiration include reactive oxygen species (ROS), highly unstable oxygen-containing molecules. Although ROS are critical for cellular signaling, an imbalance in ROS production and ROS scavenging, i.e. oxidative stress, induces oxidative damage of cellular membranes and cytotoxicity. Therefore, cells must have adequate mechanisms for scavenging mitochondrial ROS to prevent cellular damage and maintain viability.

One of the primary mechanisms for quenching mitochondrial ROS is the superoxide dismutase (SOD) metalloenzymes, which catalyze the conversion of superoxide radicals (O_2_^-^) to more stable hydrogen peroxide molecules (H_2_ O_2_) and oxygen (O_2_). There are three different isoforms of SOD (11). SOD1 is the most abundant of the isoforms and is localized to the cytosol (12). SOD3 is bound to the extracellular matrix and secreted into the extracellular space (13). SOD2 expression is restricted to the mitochondrial matrix and, therefore, is the major isoform responsible for quenching ROS produced by mitochondrial respiration (14). Thus, in the brain, SOD2 is the main isozyme involved in preserving mitochondrial function and maintaining cellular viability. Accordingly, protective antioxidant enzymes, like SOD2, may be upregulated in response to neuroinflammation in the brain during HIV infection; however, SOD2 levels in the brain during HIV infection and ART have not been determined.

In this study, we used a well-characterized SIV-infected macaque model of HIV to investigate the effects of retroviral infection on SOD2 expression in the brain, focusing on some of the most abundant cell types: neurons, astrocytes, and microglia (15-19). We also examined the effect of ART treatment on SOD2 expression in each cell type. Global *SOD2* mRNA and protein levels were evaluated in gray and white matter samples from the frontal cortex of animals in each experimental group: uninfected, untreated SIV-infected (SIV), and SIV-infected ART-treated (SIV+ART). We further explored genes associated with an increase in *SOD2* during SIV encephalitis using principal component analysis (PCA) and differential expression analysis in a targeted transcriptomic approach. PCA is a mathematical technique used to reveal variations in a dataset and emphasize any strong patterns (20). This analysis revealed that expression of *SOD2*, glial fibrillary acidic protein (*GFAP*), and Major Histocompatibility Complex I (*B2M*) were critical in distinguishing SIV from uninfected brain. These findings were validated by projecting the principal component analysis into a previously published HIV encephalitis dataset (21). We then examined changes in SOD2 expression within each cell type by co-immunolabeling SOD2 and specific cell markers (β-III Tubulin, GFAP, or CD68). Together, our data demonstrate that the expression of SOD2 by astrocytes is markedly enhanced in the brain during SIV infection; however, this neuroprotective response is not sustained at comparable high levels during ART.

## Materials and Methods

### Animal Studies

Twelve juvenile male pigtailed macaques were inoculated intravenously with both the neurovirulent clone SIV/17E-Fr and the immunosuppressive swarm SIV/DeltaB670, as previously described (22). Six animals did not receive antiretroviral treatment (denoted as the SIV group) and were euthanized before 12 weeks p.i. Six animals were infected with SIV, treated with ART beginning at day 12 p.i., and then euthanized at approximately 180 days p.i (SIV+ART). The antiretroviral therapy consisted of the nucleos(t)ide reverse transcriptase inhibitors tenofovir (9-R-2-phosphonomethoxypropyl adenine (PMPA), 20 mg/kg SID SQ, Gilead Sciences, Inc, Foster City, CA) and emtricitabine (2’,3’-dideoxy-5-fluoro-3’-thiacytidine (FTC), 40 mg/kg SID SQ, Gilead Sciences, Inc, Foster City, CA), an integrase inhibitor dolutegravir (2.5 mg/kg SID SQ, ViiV Healthcare, Research Triangle, NC), four of the animals also received the CCR5 inhibitor maraviroc (200 mg BID PO, ViiV Healthcare, Research Triangle, NC). Six additional pigtailed macaques were mock infected with 2 mL of Lactated Ringers Solution and served as untreated, virus-negative controls. Individual animal details are listed in Table 1.

**Table 1.** Animals in the study. A comprehensive list of each animal identification number, age at the time of euthanasia in years, infection status, treatment with antiretroviral therapy (ART), and the total duration of SIV infection in weeks.

| Animal Identification | Age at Euthanasia (years) | Infection Status | Treatment | Length of SIV Infection (weeks) |
| --- | --- | --- | --- | --- |
| PT247 | 3.5 | Mock | None | - |
| PT333 | 5.3 | Mock | None | - |
| PT293 | 2.3 | Mock | None | - |
| PT290 | 2.5 | Mock | None | - |
| PT332 | 5.0 | Mock | None | - |
| PT334 | 5.1 | Mock | None | - |
| PT300 | 3.2 | SIV | None | 11 |
| PT268 | 6.3 | SIV | None | 9 |
| PT203 | 9.5 | SIV | None | 9 |
| PT320 | 3.6 | SIV | None | 9 |
| PT297 | 2.5 | SIV | None | 11 |
| PT274 | 6.2 | SIV | None | 7 |
| PT394 | 3.8 | SIV | ART | 26 |
| PT400 | 4.3 | SIV | ART | 26 |
| PT401 | 3.6 | SIV | ART | 26 |
| PT390 | 4.1 | SIV | ART | 26 |
| PT391 | 3.3 | SIV | ART | 26 |
| PT392 | 3.2 | SIV | ART | 26 |

At necropsy, animals were saline-perfused and tissue samples were either immersion fixed in Streck tissue fixative (150mM 2-bromo-2-nitropane-1,3-diol, 108mM diazolinyl urea, 10mM sodium citrate dihydrate and 42mM zinc sulfate heptahydrate; Streck Laboratories, Inc, La Vista, NE) followed by paraffin embedding or snap frozen in 2-methyl butane bathed in liquid N2 then storage at −80°C. The animal procedures in this study were in accordance with the principles set forth by the Institutional Animal Care and Use Committee at Johns Hopkins University and the National Research Council’s Guide for the Care and Use of Laboratory Animals (8^th^ edition).

### RNA Isolation

Samples of gray and white matter were collected from frozen frontal cortex sections using a 3mm biopsy punch and placed into individual 2mL FastPrep Lysing Matrix D RNA tubes (116913050 MP Biomedicals, Santa Ana, CA). 1 mL of RNA STAT-60 (Amsbio LLC, Cambridge, MA) was added to each tube, and the tissue was homogenized using a Fast Prep Machine (MP Biomedicals, Santa Ana, CA) 1 x 5.0 speed for 15 seconds. Tubes remained at room temperature for 5 minutes. 200µL of chloroform was added, tubes were shaken vigorously for 15 seconds, and then left to incubate at room temperature for 2-3 minutes. Tubes were centrifuged at 14k rpm for 15 minutes at 4°C and the aqueous phase was collected into clean 1.5 mL Eppendorf tubes (Eppendorf, Hauppauge, NY). 500 µL of ice-cold isopropanol was added to each tube, then samples were vortexed and incubated at −20 °C overnight. Tubes were centrifuged at 14k rpm for 15 minutes at 4°C, then the supernatant was removed and RNA pellet was washed with ice-cold ethanol. Samples were centrifuged at 10k rpm for 5 minutes at 4°C, ethanol was then removed, and RNA pellet was left to dry at room temperature. Pellet was resuspended in RQ1 DNase solution (Promega, Madison, WI) and incubated for 1 hr at 37° with continuous agitation. DNase-free RNA was then purified using the Qiagen RNeasy Mini kit according to the package insert (Qiagen, Hilden, GE). RNA concentration was determined using a Nanodrop spectrophotometer (Thermofisher Scientific, Waltham, MA).

### Quantitative Real-Time Reverse Transcriptase Polymerase Chain Reaction

Generation of cDNA was performed using a High Capacity cDNA Reverse Transcription kit (4368814 Applied Biosystems, Foster City, CA) using 500 ng RNA per sample. Duplicate samples were run along with one sample that excluded the reverse transcriptase enzyme. qPCR was performed using TaqMan Universal Master Mix II, no UNG (4440040 ThermoFisher Scientific, Waltham, MA) according to protocol. Probes for *SOD2* (Hs00167309_m1, VIC-MGB, 4331182 ThermoFisher Scientific, Waltham, MA) and *18s* (Cy5, Integrated DNA Technologies, Coralville, IA) were used. Data were analyzed using the ΔΔCt method, where the relative expression ratio of *SOD2/18s* was calculated as 2^-ΔΔCt^ (23).

### Nanostring RNA Measurements

RNA was isolated from basal ganglia of uninfected controls (n = 10), SIV-infected animals euthanized 21 days post-infection (n = 6), and SIV-infected animals euthanized 84 days post-infection (n = 13) as previously described (24). The nanostring panel included 260 genes. All RNA measurements were normalized to the four housekeeping genes of 11 that remained the most stable across conditions (*GAPDH*, Ribosome-Like Protein L13A (*RPL13A*), Succinate Dehydrogenase A (*SDHA*), and TATA Sequence Binding Protein (*TBP*)). The limit of detection and negative controls were determined as previously described (25).

### Principal Component Analysis and Projection Analysis

Principal component analysis (PCA) of normalized Nanostring expression data was performed using the prcomp() function with centering, but no scaling, in R. One gene was removed (Microtubule-associated protein 2, *MAP2*) from this analysis because one of the uninfected controls was not evaluated for this gene.

Using weights from this PCA, data from other experiments were projected into this low-dimensional space defined by SIV infection expression change. Gene expression data from HIV infection (21) were projected into the SIV PC1 using projectR (26) (github.com/genesofeve/projectoR). Two of the four groups in the previously published National NeuroAIDS Tissue Consortium (NNTC) brain microarray (21) were included in this analysis.

### Immunoblotting

Samples of snap-frozen gray and white matter from the frontal cortex were homogenized in ice-cold RIPA buffer containing protease and phosphatase inhibitors (Sigma Aldrich, St. Louis, MO), sonicated, incubated at 4°C for 2 hours, and centrifuged to remove insoluble material. Total protein concentration was estimated using the BioRad Protein Assay Kit (BioRad Laboratories, Hercules, CA) according to the manufacturer protocol. Five micrograms of protein were loaded into 26-well 10% Bis-Tris gels (BioRad Laboratories, Hercules, CA), resolved under reducing conditions using 1 × 3-(N-morpholino)propanesulfonic acid (MOPS) buffer, and transferred to polyvinylidene fluoride (PVDF) membranes (BioRad Laboratories, Hercules, CA).

Following a one-hour blocking step in Tris-buffered saline Tween-20 (TBST, Sigma Aldrich, St. Louis, MO) containing 5% nonfat dry milk, membranes were incubated with primary antibody against SOD2 (rabbit polyclonal, 1:20,000, Abcam, Cambridge, MA) and GAPDH (rabbit polyclonal, 1:1000, Santa Cruz Biotechnology, Dallas, TX) overnight at 4°C. Membranes were subsequently washed and incubated with horseradish peroxidase-conjugated secondary antibody (goat anti-rabbit IgG, 1:20,000, Dako, Carpenteria, CA). All primary and secondary antibodies were diluted in TBST containing 5% bovine serum albumin (BSA, Sigma Aldrich, St. Louis, MO). Membranes were then developed with chemiluminescent substrate (SuperSignal West Dura, ThermoFisher Scientific, Waltham, MA) and exposed to radiography film, which was digitally scanned. Densitometry was performed using ImageJ software. SOD2 protein levels were normalized to GAPDH and reported as relative fold change (compared to uninfected controls).

### Immunohistochemistry

Immunohistochemistry was performed on Streck-fixed, paraffin-embedded sections of frontal cortex using a Leica Biosystems Bond RX^m^ automated staining system (Leica Biosystems, Wetzler, Germany). Tissues were deparaffinized with Bond Dewax Solution (AR9222, Leica Biosystems, Wetzler, Germany) 3× 1 min then washed with 70% alcohol 3x followed by Bond Wash solution (AR9590, Leica Biosystems, Wetzler, Germany) 3x. Antigen retrieval was done by incubation with Bond Epitope Retrieval solution 2 (AR9640, Leica Biosystems, Wetzler, Germany) for 20 minutes and then sections were then washed with Bond Wash solution 3x. Immunostaining was performed using reagents provided by the Bond Polymer Refine Detection kit (DS9800, Leica Biosystems, Wetzler, Germany). Sections were blocked with Peroxide Block solution (Leica Biosystems, Wetzler, Germany) for 5 minutes and washed 3x with Bond Wash solution. Sections were then incubated with the primary antibody (Rabbit anti-SOD2, 1:2500, ab13534, Abcam, Cambridge, MA) diluted in Bond Primary Antibody Diluent (AR9352, Leica Biosystems, Wetzler, Germany), for 15 minutes. Sections were washed 3x with Bond Wash solution and incubated in Post Primary solution for 8 minutes. Sections were washed 3× 2 minutes and incubated in Polymer solution for 8 minutes. Bond Wash solution was applied 2× 2 minute incubations followed by a wash with deionized water. Sections were washed once with Mixed DAB Refine Solution (Leica Biosystems, Wetzler, Germany) and then incubated in Mixed DAB Refine Solution for 10 minutes. Slides were washed 3x with deionized water and counterstained with Hematoxylin (Leica Biosystems, Wetzler, Germany) for 5 minutes. Sections were then washed once with deionized water, once with Bond Wash solution, and once again with deionized water. After staining was finished, the slides were dehydrated with increasing concentrations of ethanol (70% ethanol for 2×1min, 90% ethanol for 2×1min, 100% ethanol for 2×1min), cleared in Histoclear (HS-200, National Diagnostics, Atlanta, GA) for 2×5min, and coverslipped with Permount mounting medium (Fisher Scientific, Pittsburgh, PA). Image acquisition and quantification of positive SOD2 immunostaining was performed using a Nikon DS-Ri1 color camera mounted on a Nikon Eclipse 90i microscope and NIS Elements software (vAR710, Nikon, Melville, NY). A composite image of a slice through the frontal cortex was created by aligning serial images of contiguous 20× fields from the cortical surface extending through underlying white matter tracts. Quantification of SOD2 in gray and white matter was performed by manually tracing regions of interest (ROI), using digital thresholding to define pixels positive for SOD2 stain, and calculating the fraction of the total ROI area occupied by positive staining.

### Immunofluorescence Labeling and Confocal Microscopy

Immunofluorescence labeling was performed on Streck-fixed paraffin-embedded sections of frontal cortex using a Leica Biosystems Bond RX^m^ automated staining system (Leica Biosystems, Wetzler, Germany). Tissue deparaffinization and antigen retrieval were performed as indicated above for Immunohistochemistry. Following antigen retrieval, sections were washed with Bond Wash solution 2x 10s and blocked with a phosphate-buffered saline (PBS) solution containing 5% normal goat serum (5425S, Cell Signaling, Danvers, MA) and 0.5% TritonX-100 (T8787, Sigma Aldrich, St. Louis, MO) for 15 minutes. Slides were washed with Bond Wash solution 3x 30s then incubated with a rabbit anti-SOD2 primary antibody (ab13534, abcam, Cambridge, MA) 1:500 in Bond Primary Antibody Diluent for 1 hour. Slides were washed with Bond Wash solution 3× 3 minutes then incubated with a goat anti-rabbit IgG (H+L) AlexaFluor 633-conjugated secondary antibody (A-21070, ThermoFisher Scientific, Waltham, MA) 1:500 in PBS containing 3% goat serum, 0.1% TritonX-100 for 1 hour. Slides were then washed with Bond Wash solution 3 × 5 minutes and blocked again with PBS (0.5% goat serum, 0.5% TritonX-100) for 15 minutes. Sections were washed with Bond Wash buffer 3 x 30s and then incubated with either mouse anti-CD68 (M0814, 1:100, Dako, Carpenteria, CA), mouse anti-βIII-Tubulin (G7121, Promega, Madison, WI), mouse anti-GFAP (PA0026, Leica Biosystems, Wetzler, Germany), or mouse anti-Factor VIII (ab41187, Abcam, Cambridge, MA) diluted in Bond Primary Antibody Diluent for 1 hour. Sections were washed Bond Wash solution for 3 × 3 minutes and then incubated with a goat anti-mouse IgG (H+L) AlexaFluor 488-conjugated secondary antibody (A-1101, ThermoFisher Scientific, Waltham, MA) 1:500 in PBS containing 3% goat serum, 0.1% TritonX-100 for 1 hour. Slides were washed with Bond Wash solution 3 × 5 minutes and then wet mounted with ProLong Gold Antifade reagent with DAPI (P36931, Invitrogen, Carlsbad, CA) and allowed to dry overnight in the dark. Co-localization of the resulting red (SOD2) and green (GFAP, βIII-Tubulin, CD68, or Factor VIII) fluorescent labeling was visualized using a Nikon C1 confocal laser microscope system mounted on a Nikon Eclipse TE2000-E microscope in conjunction with EZ-C1 software (v3.4). Fluorescently labeled slides were compared to slides that underwent the same protocol above with isotype control antibodies used in place of the primary antibodies (rabbit IgG1 2 µg/mL for SOD2, mouse IgG1 0.5 µg/mL for GFAP, mouse IgG1 10 µg/mL for βIII-Tubulin, and mouse IgG1 2 µg/mL for CD68).

### Statistics

All statistical inferences were calculated using nonparametric methods and GraphPad Prism Software (v5.0d, GraphPad Software, San Diego, CA). Group comparisons were performed using a one way-ANOVA followed by a Benjamini-Hochberg post-hoc test. For the Nanostring and microarray datasets, Mann-Whitney comparisons were done in R without correction. For all analyses, statistical significance was assumed when the *p* value was less than 0.05.

### Data Availability

The data that support the findings of this study are available from the corresponding author upon reasonable request.

## Results

### SOD2 Expression Was Upregulated in the Brain of Untreated SIV-Infected Animals and Normalized with ART

We measured *SOD2* mRNA levels using reverse transcriptase quantitative polymerase chain reaction (qRT-PCR) in samples of frontal cortex from uninfected and SIV-infected pigtailed macaques. An approximately 10-fold increase in *SOD2* levels was observed in gray matter samples from SIV-infected animals relative to uninfected animals (Figure 1A). *SOD2* mRNA levels were dramatically elevated by nearly 100-fold in white matter samples from SIV-infected pigtailed macaques relative to uninfected controls (Figure 1B). Our findings support previously published data that show increased *SOD2* levels in primary cultured cells infected with HIV and in dorsal root ganglia during acute SIV infection (8, 27). In this report, *SOD2* mRNA levels in SIV+ART animals were intermediate between levels present in SIV and uninfected animals in both gray and white matter. Together, these data indicate that *SOD2* is upregulated in gray and white matter during SIV-infection, and that this upregulation is attenuated by ART.

**Figure 1.**
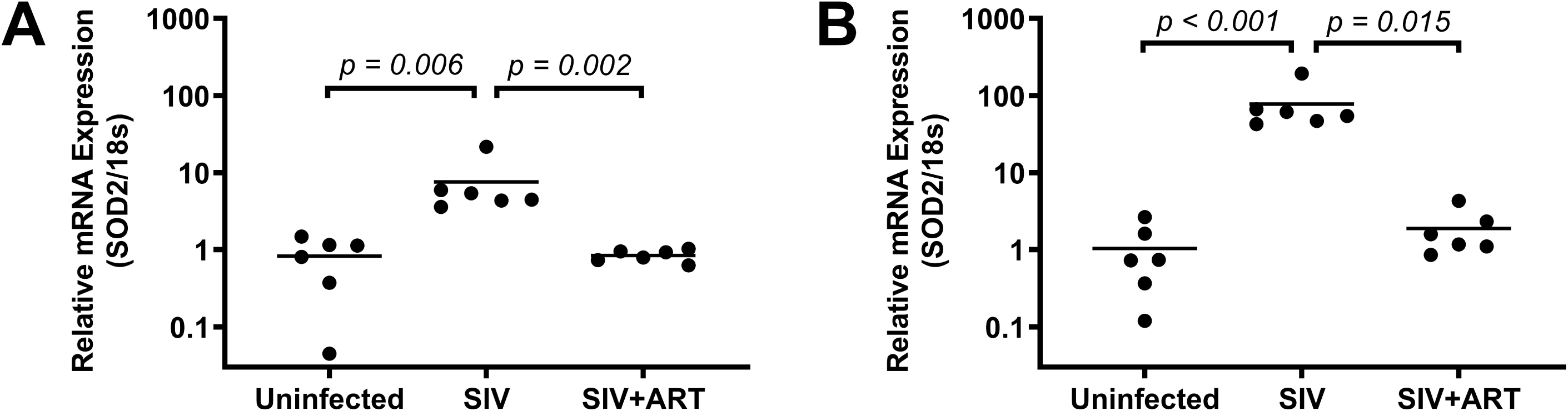
Summary data indicating relative mRNA levels of superoxide dismutase 2 (SOD2) to 18s in cortical gray matter (A) and white matter (B) from uninfected, SIV-infected, and antiretroviral therapy (ART)-treated SIV-infected animals (SIV+ART). Data were normalized to mean value for uninfected. Bars represent median values; n = 6 animals per group.

Expanding on the global *SOD2* mRNA levels measured by qPCR, we validated these findings using a Nanostring dataset run previously (24). In this dataset, *SOD2* mRNA counts were significantly increased after SIV infection (Figure 2A; *p* < 0.0001). Similarly, *GFAP* mRNA was significantly increased in this group compared to uninfected controls in this dataset (Figure 2B; *p* < 0.0005). The increase in *SOD2* mRNA counts was also significantly correlated with the increase observed in *GFAP* mRNA counts (Figure 2C; Spearman *R* = 0.79, *p* = 0.003). These data suggest that increased transcription of *SOD2* and *GFAP* are moving in lockstep during SIV infection.

**Figure 2.**
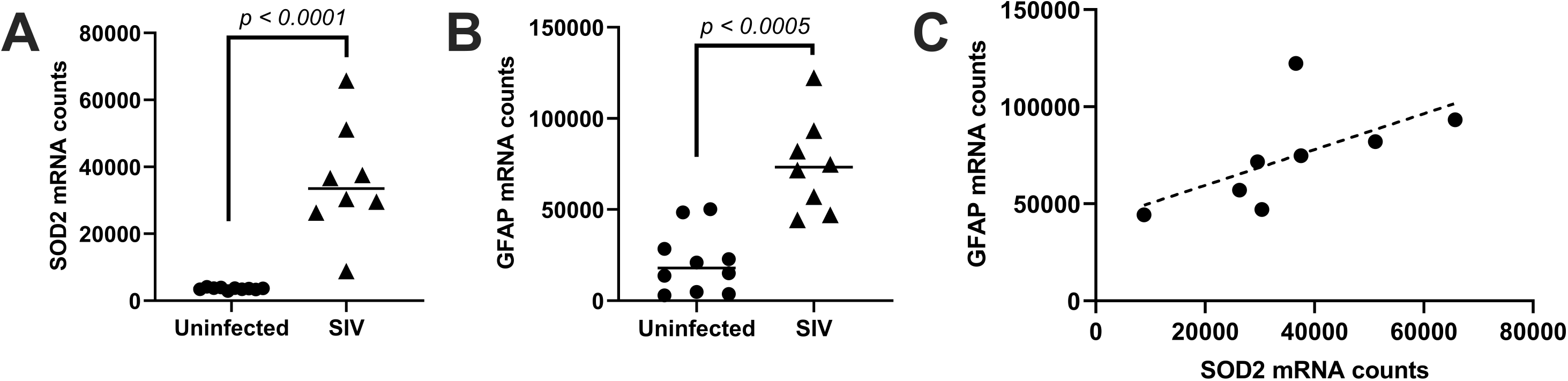
Summary data indicating mRNA counts for (A) superoxide dismutase 2 (SOD2) and (B) glial fibrillary acidic protein (GFAP) for uninfected controls and SIV-infected animals. C) Correlation of SOD2 and GFAP mRNA counts in SIV-infected animals. Bars represent median value; n = 8 animals per group.

To further explore this relationship, principal component analysis (PCA) was performed on this dataset. Principal Component 1 (PC1) accounted for 79.1% of the variance between samples (Figure 3). The SIV group clusters away from the uninfected controls along PC1. The weights for individual genes in this PC1 indicate the extent to which each gene contributes to this major trajectory of expression change during SIV infection. Among the individual genes driving PC1, *SOD2* and *GFAP* were among the top three differentially expressed genes, though Major Histocompatibility Complex I (MHC I, *B2M*) was the top differentially expressed gene (Supplemental Table 1). These data indicate that *SOD2* and *GFAP* are specifically upregulated in the brain during SIV encephalitis.

**Figure 3.**
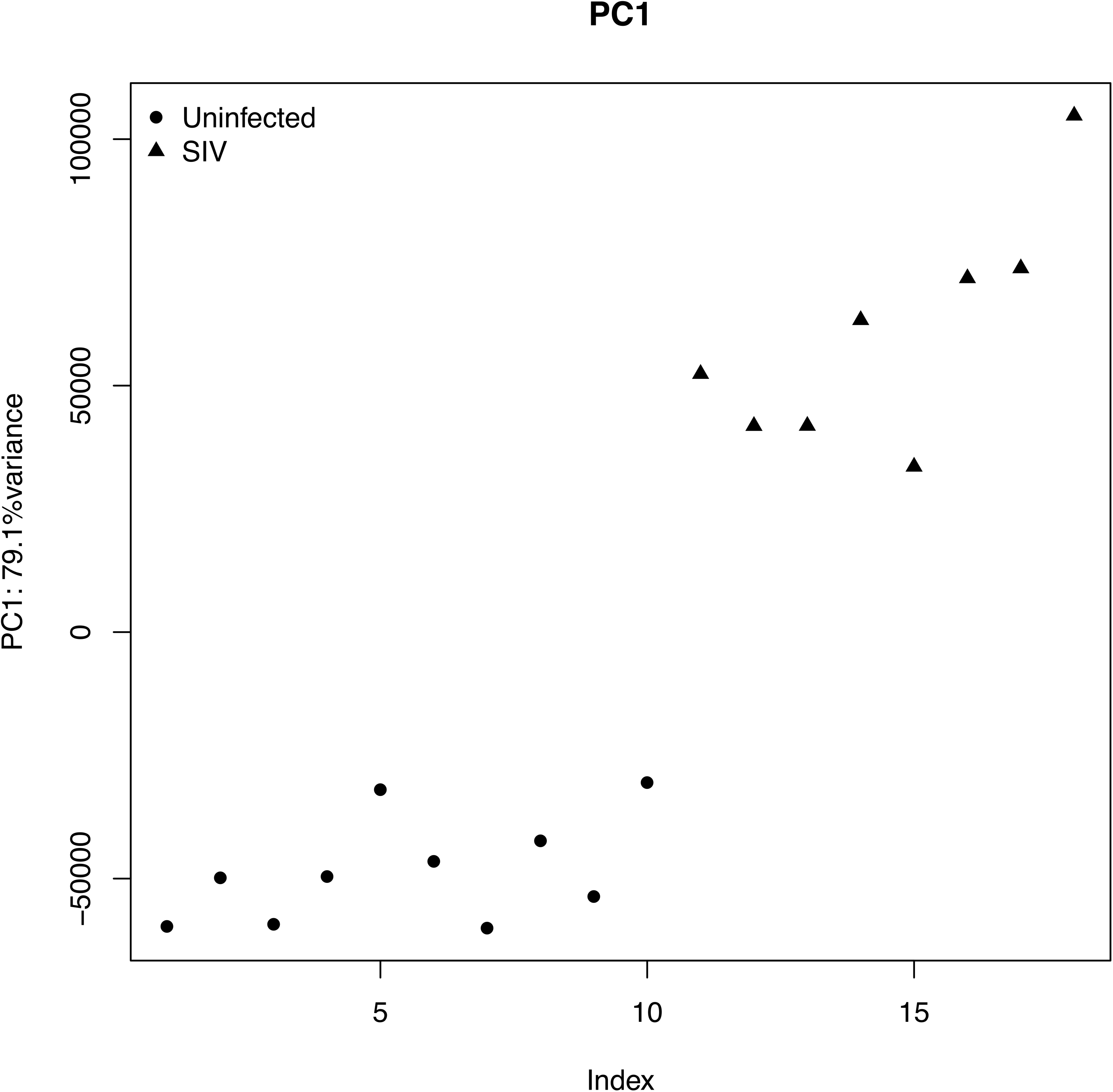
Principal component analysis of Nanostring data from uninfected and SIV-infected basal ganglia. Principal Component 1 (PC1), which is responsible for 79.1% of the variance in the data, clearly separates uninfected (n = 10, •) from SIV-infected (n = 8, ▲) animals. Genes defining this separation include superoxide dismutase 2 (*SOD2*), glial fibrillary acidic protein (*GFAP*), and Major Histocompatibility Complex I (*B2M*).

### SOD2 expression in HIV

To determine whether these changes also occur in individuals infected with HIV, we utilized the previously published National NeuroAIDS Tissue Consortium brain microarray data (21). *SOD2* (Figure 4A; *p* = 0.001) and *GFAP* (Figure 4B; *p* = 0.0002) expression were significantly increased in individuals with HIV encephalitis (HIVE) compared to uninfected individuals. Although *SOD2* and *GFAP* mRNA expression are not well correlated (data not shown; Spearman R = 0.25), there is a clear separation between the uninfected controls and the HIVE-diagnosed individuals when plotting *SOD2* and *GFAP* mRNA levels (Supplemental Figure 1). This suggests that although not directly correlated, increased *SOD2* and *GFAP* are defining characteristics of HIVE, similar to SIV-infected animals.

**Figure 4.**
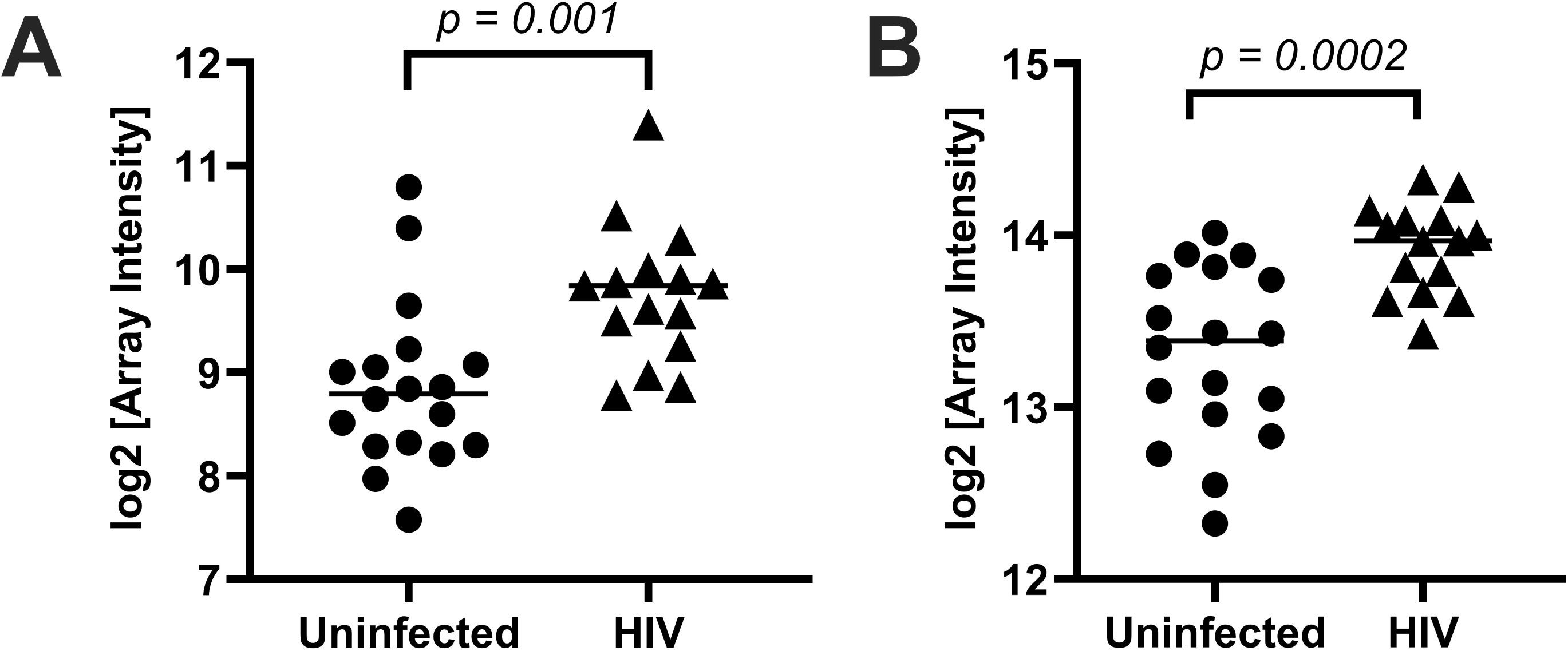
Summary data indicating the expression (log2 [Array Intensity]) for (A) superoxide dismutase 2 (SOD2) and (B) glial fibrillary acidic protein (GFAP) for uninfected individuals and individuals with HIV encephalitis. Bars represent median value; n = 15-18 people per group.

While this shows similarities in transcriptional changes in SIV and HIV, this method is restricted to interrogating relatively few genes. To be able to determine transcriptional program changes, global analyses must be utilized. The SIV transcriptional analysis was based on a Nanostring panel of 250 genes. To relate the dynamics of this SIV infection model to HIV infection, we projected gene expression data from postmortem brain tissue of uninfected individuals and patients diagnosed with HIVE (19) into the transcriptional space defined by the individual gene weights from PC1 using projectR (20). This projection produced a clear separation between the HIVE samples and the uninfected control samples (Figure 5), demonstrating similarity between the transcriptional changes induced in SIV and HIV. The genes primarily driving this separation are again *B2M, SOD2,* and *GFAP* (Supplemental Table 1), indicating that expression changes in *SOD2* and *GFAP* specifically are characteristic of both SIV and HIV.

**Figure 5.**
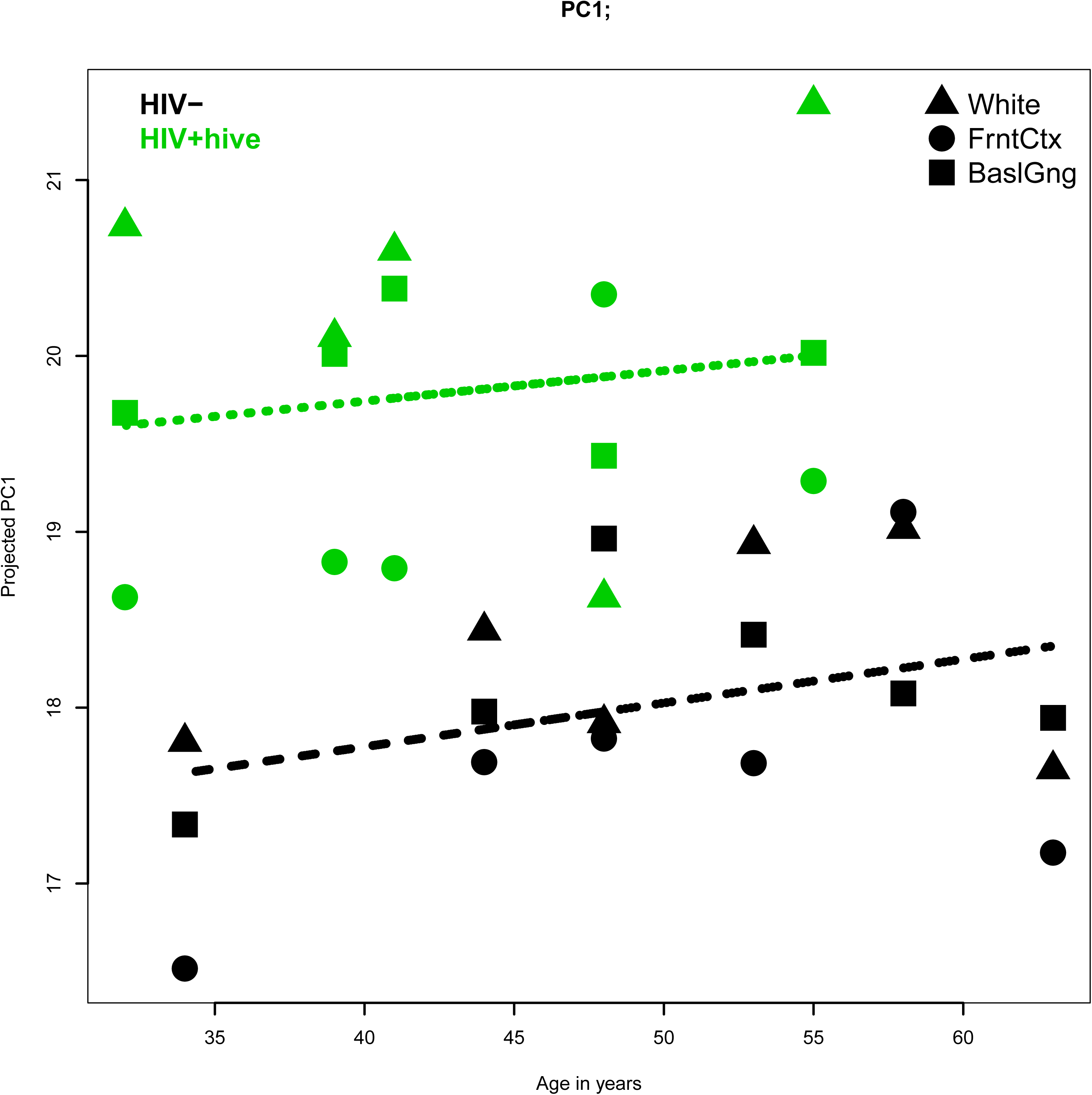
Projection of Principal Component 1 (PC1) from Nanostring transcriptomic changes in uninfected and SIV-infected macaque basal ganglia into previously published HIV encephalitis (HIVE) microarray data separates HIVE from uninfected individuals (uninfected n = 6, HIVE n = 5; ▲ = white matter, l1 = basal ganglia, • = frontal cortex).

### SOD2 Protein Was Enhanced in the Brain of Untreated SIV-Infected Animals and Normalized with ART

We next examined global SOD2 protein levels in the brain of SIV-infected and uninfected animals. Immunoblotting for SOD2 was performed to compare the SOD2 protein levels between the treatment groups. Similar to SOD2 RNA increases, SOD2 protein was found to be increased in SIV-infected gray and white matter when compared with uninfected animals (Figures 6A and 6B). SOD2 protein in gray and white matter samples from SIV+ART animals was similar to uninfected controls.

**Figure 6.**
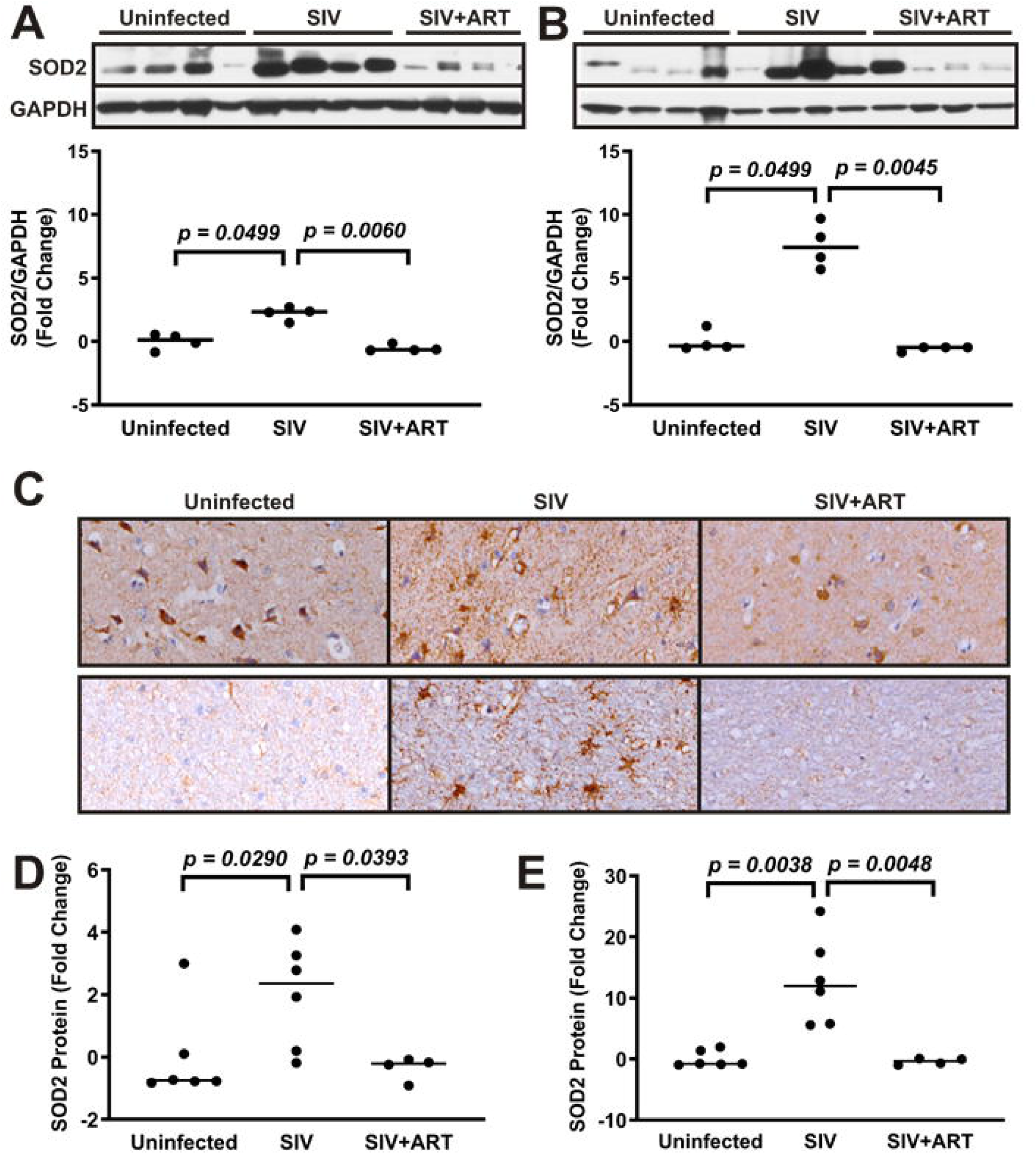
A and B) Representative immunoblot for superoxide dismutase 2 (SOD2, 25 kDa) and GAPDH (37 kDa) in uninfected, SIV-infected, and antiretroviral therapy (ART)-treated SIV-infected animals (SIV+ART) frontal cortex gray matter (A) and white matter (B). SOD2 protein density was normalized to GAPDH for each sample and represented as Fold Change from Uninfected. Bars indicate median values; n = 4 animals per group. C) Representative images of gray matter (top) and white matter (bottom) indicating immunostaining for SOD2. D and E) The fraction of area positively stained for SOD2 in gray matter (D) and white matter (E) was calculated and represented as Fold Change from the uninfected mean. Bars represent median value; n = 4-6 animals per group.

To investigate cellular changes in SOD2 protein, we performed immunohistochemical staining for SOD2 in sections of frontal cortex. In uninfected control animals, SOD2 staining primarily was localized to round to polygonal cells in gray matter (likely neuronal cell bodies) and was nearly absent in white matter (Figure 6C). Brain tissue from untreated SIV-infected animals exhibited abundant SOD2 staining, which displayed a stellate pattern in both gray and white matter, more consistent with glial cells (astrocytes and/or microglia). Quantitation of SOD2 staining in these sections using digital image analysis revealed that SOD2 protein levels were more than 10x higher in gray matter and over 50x higher in white matter of SIV-infected animals relative to uninfected controls (Figures 6D-E). Consistent with our findings by Western blot, SOD2 immunostaining was normalized in SIV+ART animals. Together with our SOD2 mRNA findings, these findings indicate that SOD2 is upregulated in glial cells of the brain during SIV infection, but normalized in animals receiving ART.

### SOD2 Was Increased in Astrocytes but Reduced in Neurons of SIV-Infected Animals, and These Changes are Normalized with ART

Based on the stellate, branching immunohistochemical staining pattern observed in SIV-infected animals, we hypothesized that SOD2 was primarily upregulated in astrocytes in the brain during SIV infection. To determine cell-specificity of SOD2 upregulation, we performed double immunofluorescent labeling of brain sections and imaged these using confocal microscopy.

First, we co-immunolabeled for SOD2 and the activated astrocytic marker glial fibrillary acidic protein (GFAP). GFAP immunolabeling was detected in classic stellate patterns in both gray and white matter of uninfected and SIV-infected animals. SOD2 immunolabeling was much more apparent in gray and white matter of SIV-infected animals relative to uninfected controls and also displayed a punctate pattern within the brain parenchyma (Figures 7A and 7B), consistent with previous reports (28). In SIV-infected animals, there was abundant co-localization of SOD2 and GFAP immunostaining, demonstrating that SOD2 was increased in astrocytes in these animals.

**Figure 7.**
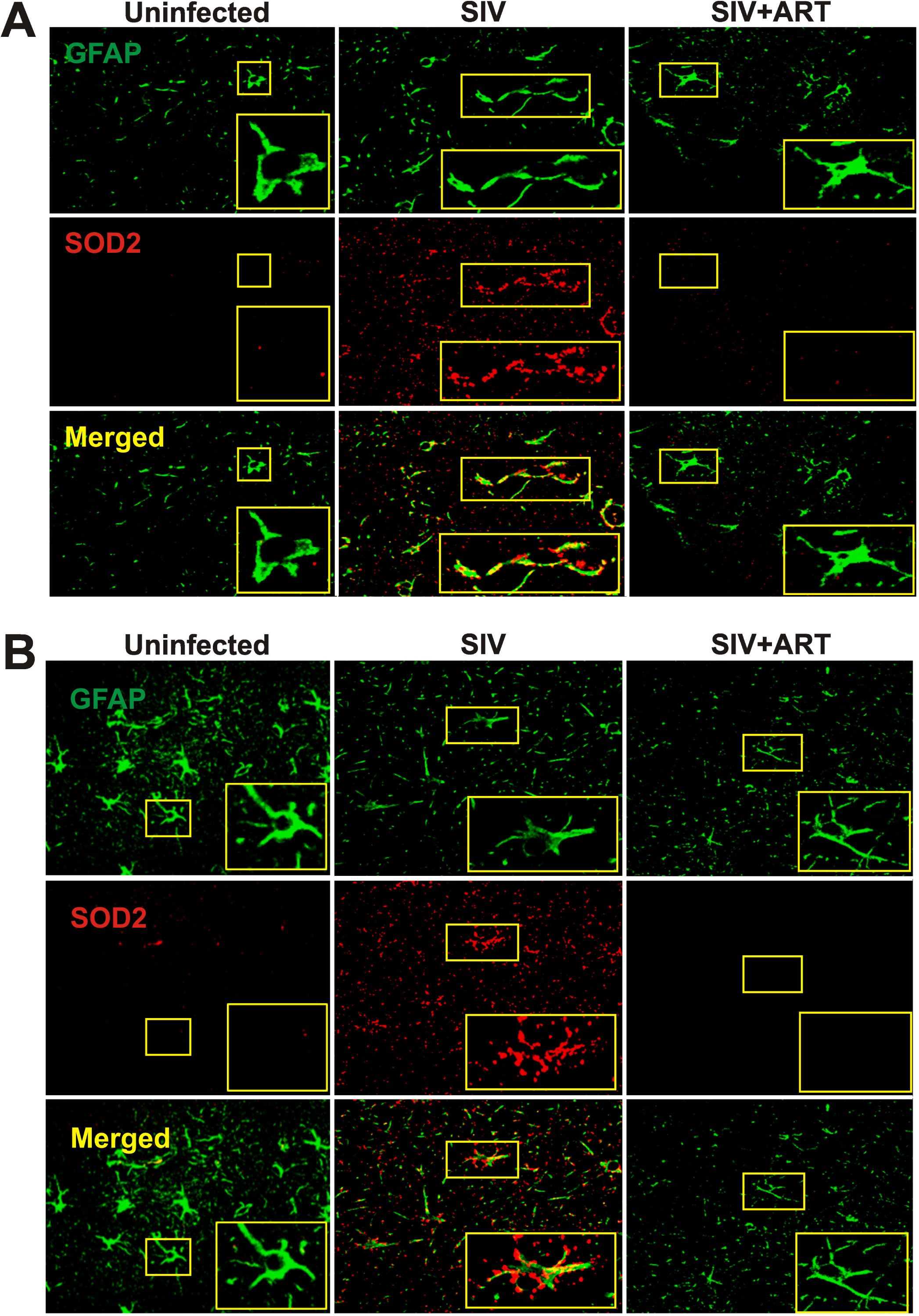
Representative confocal images of frontal cortex gray matter (A) and white matter (B) immunofluorescently co-labeled for the astrocytic marker glial fibrillary acidic protein (GFAP, green) and superoxide dismutase 2 (SOD2, red). Merged images indicate areas of SOD2 co-localization with GFAP (yellow).

SOD2 levels within neurons and microglia were also evaluated. Immunolabeling for SOD2 and a neuron-specific protein, β-III tubulin, revealed low levels of SOD2 that co-localized with labeling for β-III tubulin in uninfected controls. In SIV-infected animals, SOD2 immunolabeling was not detected above background levels within neurons; however, strong punctate labeling was observed surrounding the neurons and within the parenchyma in stellate patterns (Figure 8). Co-association of SOD2 with CNS macrophages, including parenchymal microglia and perivascular macrophages, was determined by immunolabeling for SOD2 and CD68, a protein that is upregulated in activated macrophages and microglia (29). SOD2 immunolabeling co-localized with perivascular macrophages in uninfected animals (Figure 9). In SIV-infected animals, both SOD2 and CD68 immunolabeling increased in perivascular and parenchymal clusters of glial cells; however, there was little overlap. These data indicate that SOD2 is reduced in neurons but unchanged in microglia during SIV infection.

**Figure 8.**
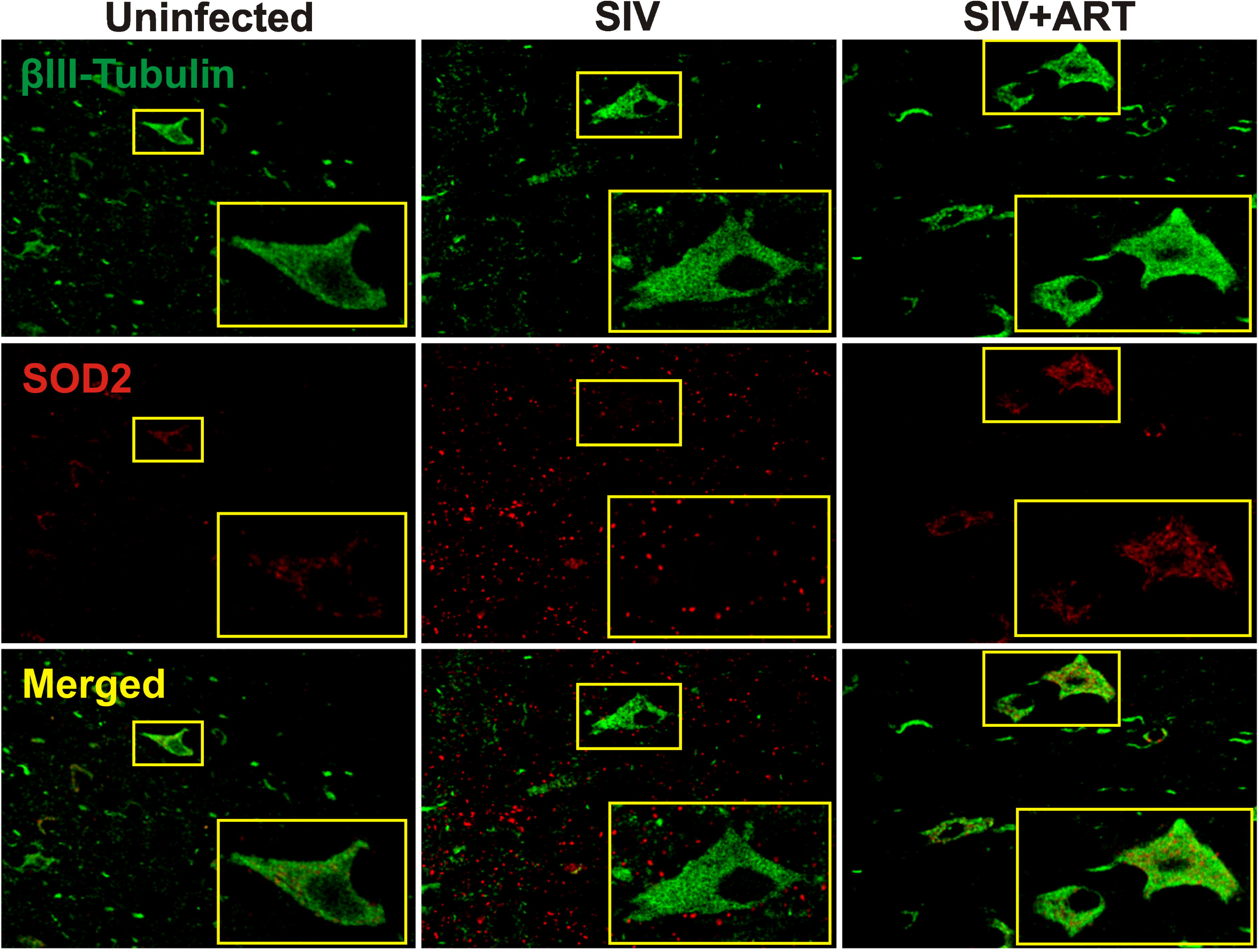
Representative confocal images of frontal cortex gray matter immunofluorescently co-labeled for the neuronal marker βIII-Tubulin (green) and superoxide dismutase 2 (SOD2, red). Merged images indicate areas of SOD2 co-localization with βIII-Tubulin (yellow).

**Figure 9.**
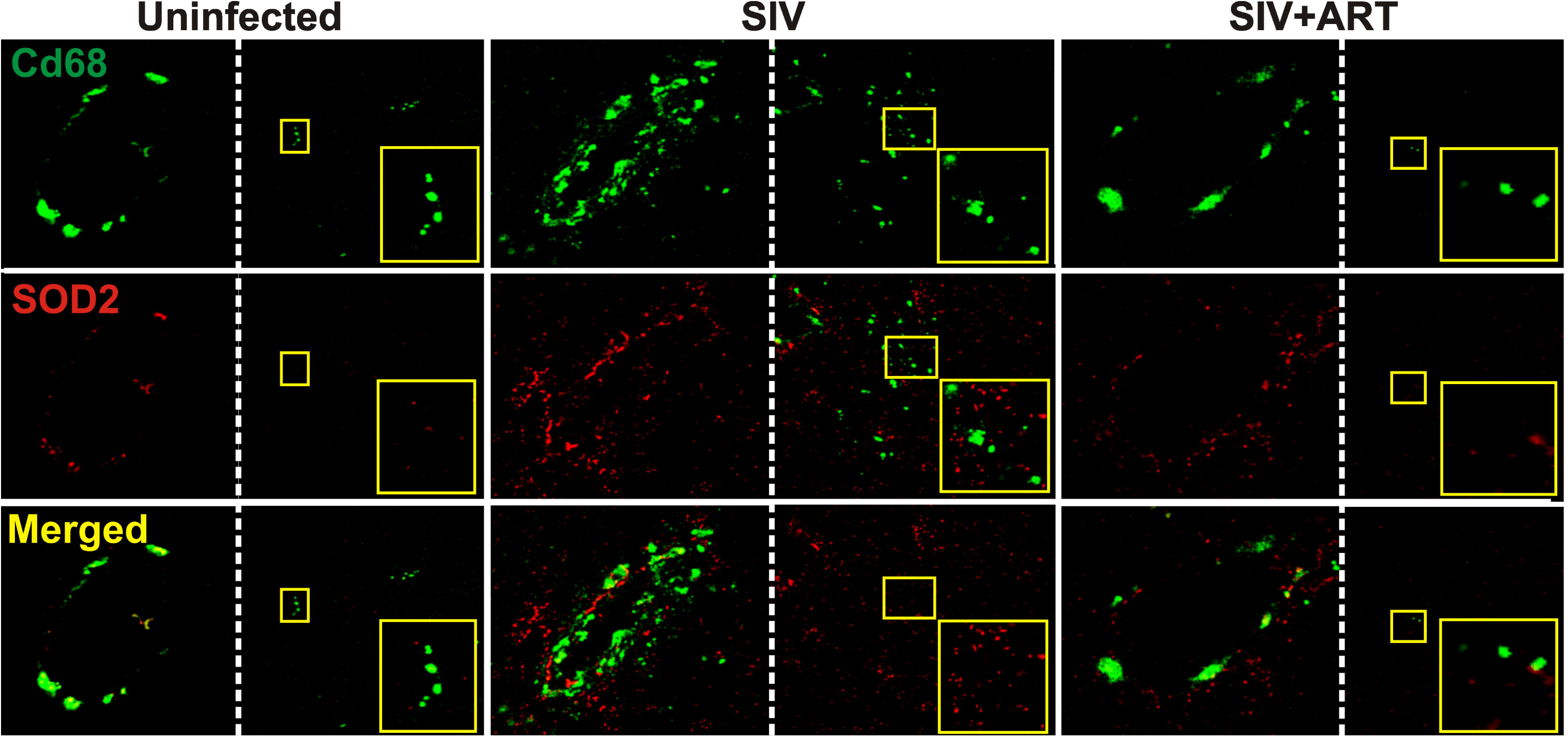
Representative confocal images of frontal cortex immunofluorescently co-labeled for the microglial marker CD68 (green) and superoxide dismutase 2 (SOD2, red). CD68+ microglia were prominent surrounding blood vessels (left images) but also found in clusters within the brain tissue (right images). Merged images indicate areas of SOD2 co-localization with CD68 (yellow).

In SIV+ART animals, SOD2 labeling was minimal with little to no co-localization of SOD2 and GFAP. SOD2 and β-III tubulin co-localization in SIV+ART animals matched the pattern present in uninfected animals. The extent of co-localization of SOD2 and CD68 in SIV+ART animals also appeared similar to that observed in uninfected animals. Together, our findings demonstrated that ART reduced SOD2 protein levels from the peaks seen in untreated SIV animals within each cell type to levels similar to those found in uninfected animals.

## Discussion

Our data indicate that SOD2 is elevated in the brain during SIV infection primarily due to upregulation by astrocytes, and we confirmed that similar changes occur in HIV patients with encephalitis. The primary function of astrocytes is to support and preserve neuronal function; therefore, disruption of normal astrocytic function can lead to neurotoxic conditions. Astrocyte dysfunction has been implicated as a critical factor in the development of neurocognitive deficits during HIV infection as well as with other neurodegenerative diseases, such as Alzheimer’s Disease and Huntington’s Disease (30). Widespread astrocytosis is a classic neuropathologic finding in our SIV model (31, 32) as well as HIV-infected humans (33). Although astrocytes can be infected by HIV (34), they lack the ability to generate replication-competent virus (35-37). Interestingly, the frequency of HIV infection of astrocytes has been shown to be positively correlated with the severity of neuropathologic conditions in human patients (38), suggesting that astrocytes may produce neurotoxic viral proteins. Furthermore, using sensitive DNAscope approaches, a recent study has indicated that astrocytes do not harbor HIV DNA after long-term ART in contrast with microglia (39). Interestingly, data presented here suggest that SOD2 is upregulated in the brain of HIV patients when compared to uninfected individuals. Independent of HIV infection of astrocytes, it is possible that SOD2 upregulation helps to preserve astrocytic function during CNS HIV infection, thus delaying the development of neurocognitive deficits. Studies examining the effects of manipulating SOD2 expression or function in SIV infected macaques could clarify this potential role of SOD2.

Although the mechanisms underlying the upregulation of SOD2 by astrocytes in response to SIV infection are unknown, the primary inciting cause is likely the pro-neuroinflammatory state of the brain during retroviral infection. The SIV/pigtailed macaque model has been shown to develop significant neuroinflammation during infection, similar to what is observed in human neuroAIDS patients (8, 17, 40). The data presented here illustrate that the SIV/pigtailed macaque model is a highly relevant model to study changes in central nervous system gene expression that accompany HIV brain infection (Figure 5). SOD2 transcription is induced by many pro-inflammatory cytokines, including IL-1α, IL-1β, and TNFα (41), therefore, generalized neuroinflammation has been proposed as the primary stimulus for SOD2 upregulation in the brain. However, *in vitro* experiments show that the addition of a classic inflammatory molecule, lipopolysaccharide, alone does not upregulate SOD2 in primary rat astrocytes (42), suggesting that a virus-specific mechanism may be required for astrocytic upregulation of SOD2. The HIV-secreted protein, Tat, has been shown to induce gene expression and increase oxidative stress in astrocytes (43). Another *in vitro* study demonstrated that the HIV envelope glycoprotein, gp120, activates NF-kB p65 and increases SOD2 expression in human primary astrocytes (27). Whether SIV-specific molecules like Tat and gp120 induce a compensatory elevation in antioxidant molecules or directly alter SOD2 gene regulation in our model remains to be determined.

Our fluorescent co-localization studies showed that SOD2 was not upregulated in neurons of animals infected with SIV (Figure 8). Although neurons are not infected with SIV/HIV, they are susceptible to damage by neurotoxic viral proteins and inflammatory mediators released by infected and/or reactive glial cells (44). Compared to astrocytes, neurons have a reduced ability to upregulate SOD2 during inflammation and oxidative insult due to lower basal levels of NFkB p65 (27). Therefore, it is possible that, during SIV-infection, lack of neuronal SOD2 upregulation contributes due to progressive mitochondrial damage and neuronal dysfunction.

Interestingly, despite extensive literature describing productive infection and activation of microglia during SIV and HIV infection (32, 45, 46) as well as ongoing microglial expression of CSF1R despite suppressive ART (25), our findings indicate that microglial SOD2 remained unchanged during SIV infection and SIV+ART (Figure 9). Models using an intracerebral injection of lipopolysaccharide have shown that microglia upregulate SOD2 during generalized neuroinflammation (42). However, reports examining HIV-infected microglia indicate that cytosolic SOD1 is significantly upregulated (47), suggesting that the state of oxidative stress may be higher in the cytosol than in the mitochondria of microglia during virus-induced neuroinflammation. Follow-up immunolocalization studies are needed to determine if SOD1 is upregulated in microglia in our SIV model.

HIV-infected patients still develop neurocognitive disorders and have elevated markers of oxidative stress in the brain despite ART (48). Although there is evidence that ART reduces neuroinflammation and resolves reactive astrogliosis (49), persistent ROS accumulation due to persistent low-level inflammation and chronic administration of neurotoxic ART drugs may contribute to neuronal dysfunction and cognitive deficits (49, 50). In the current study, total brain SOD2 mRNA and protein levels, as well as neuronal and astrocytic SOD2 levels, were lower in SIV+ART animals. This finding was likely related to early, rapid suppression of SIV in the plasma and CSF and the relatively short duration of the study (3 months). Also, three of the four ART drugs used in this study (tenofovir, emtricitabine, and maraviroc), show low levels of neurotoxicity *in vitro* (50). Dolutegravir has been found to be neurotoxic at higher doses *in vitro* and *in vivo* (51, 52). It is also possible that ART drugs have off-target effects that impair the compensatory upregulation SOD2. Mechanistic *in vitro* studies will be necessary to understand the impact of specific ART compounds on SOD2 regulation in neurons and astrocytes.

In conclusion, our study found that SOD2 is enhanced primarily in astrocytes during SIV infection, and this change is not maintained at comparable high levels with ART. The upregulation of SOD2 expression by astrocytes is likely a neuroprotective mechanism in response to viral-induced neuroinflammation. Therefore, developing therapies that are aimed at enhancing SOD2 activity in the brain may be useful to mitigate neuronal damage and dysfunction associated with HIV infection and may reduce the incidence or severity of HAND.

## Author Roles

MNS performed experiments, analyzed data, prepared figures, and wrote the initial draft and edited the manuscript. SAB, YJJ, CVS, ACK, and CC performed experiments and provided comments on the manuscript. SEQ provided data and comments on the manuscript. GK and NP provided comments on the manuscript. LMM and JLM conceived the study and edited the manuscript.

## Acknowledgements

National Institutes of Health (R01NS089482 to JLM, U42OD013117 to JLM, R01HL136918 to NP, and R01HL063030 to NP), the Boehringer Ingelheim Veterinary Scholars Program (MNS), and a T32 training grant T32AG058527 (GK) provided support for these studies. The JHU Magic-That-Matters Fund (NP) also provided support for these studies. We thank Gilead Sciences, Inc. for their generous donation of TFV and FTC. We thank ViiV Healthcare for their kind donation of dolutegravir and maraviroc. We acknowledge the contribution of the JHU Retrovirus Lab team members in assisting the performance and analysis of these studies.

## References

1. de Mendoza C. UNAIDS Update Global HIV Numbers. AIDS Rev 2019;21:170–1

2. Yuan NY, Kaul M. Beneficial and Adverse Effects of cART Affect Neurocognitive Function in HIV-1 Infection: Balancing Viral Suppression against Neuronal Stress and Injury. J Neuroimmune Pharmacol 2019

3. Heaton RK, Franklin DR, Ellis RJ, et al. HIV-associated neurocognitive disorders before and during the era of combination antiretroviral therapy: differences in rates, nature, and predictors. J Neurovirol 2011;17:3–16

4. Saylor D, Dickens AM, Sacktor N, et al. HIV-associated neurocognitive disorder--pathogenesis and prospects for treatment. Nat Rev Neurol 2016;12:234–48

5. Burte F, Carelli V, Chinnery PF, et al. Disturbed mitochondrial dynamics and neurodegenerative disorders. Nat Rev Neurol 2015;11:11–24

6. Rozzi SJ, Avdoshina V, Fields JA, et al. Human Immunodeficiency Virus Promotes Mitochondrial Toxicity. Neurotox Res 2017;32:723–33

7. Lehmann HC, Chen W, Borzan J, et al. Mitochondrial dysfunction in distal axons contributes to human immunodeficiency virus sensory neuropathy. Ann Neurol 2011;69:100–10

8. Mangus LM, Weinberg RL, Knight AC, et al. SIV-Induced Immune Activation and Metabolic Alterations in the Dorsal Root Ganglia During Acute Infection. J Neuropathol Exp Neurol 2019;78:78–87

9. Picard M, McEwen BS. Mitochondria impact brain function and cognition. Proc Natl Acad Sci U S A 2014;111:7–8

10. Silver I, Erecinska M. Oxygen and ion concentrations in normoxic and hypoxic brain cells. Adv Exp Med Biol 1998;454:7–16

11. Fridovich I. Superoxide dismutases. Annu Rev Biochem 1975;44:147–59

12. Crapo JD, Oury T, Rabouille C, et al. Copper,zinc superoxide dismutase is primarily a cytosolic protein in human cells. Proc Natl Acad Sci U S A 1992;89:10405–9

13. Marklund SL, Holme E, Hellner L. Superoxide dismutase in extracellular fluids. Clin Chim Acta 1982;126:41–51

14. Sun J, Ren X, Simpkins JW. Sequential Upregulation of Superoxide Dismutase 2 and Heme Oxygenase 1 by tert-Butylhydroquinone Protects Mitochondria during Oxidative Stress. Mol Pharmacol 2015;88:437–49

15. Gama L, Abreu C, Shirk EN, et al. SIV Latency in Macrophages in the CNS. Curr Top Microbiol Immunol 2018;417:111–30

16. Mangus LM, Dorsey JL, Laast VA, et al. Neuroinflammation and virus replication in the spinal cord of simian immunodeficiency virus-infected macaques. J Neuropathol Exp Neurol 2015;74:38–47

17. Laast VA, Shim B, Johanek LM, et al. Macrophage-mediated dorsal root ganglion damage precedes altered nerve conduction in SIV-infected macaques. Am J Pathol 2011;179:2337–45

18. Laast VA, Pardo CA, Tarwater PM, et al. Pathogenesis of simian immunodeficiency virus-induced alterations in macaque trigeminal ganglia. J Neuropathol Exp Neurol 2007;66:26–34

19. Beck SE, Kelly KM, Queen SE, et al. Macaque species susceptibility to simian immunodeficiency virus: increased incidence of SIV central nervous system disease in pigtailed macaques versus rhesus macaques. J Neurovirol 2015;21:148–58

20. Hess AS, Hess JR. Principal component analysis. Transfusion 2018;58:1580–2

21. Gelman BB, Chen T, Lisinicchia JG, et al. The National NeuroAIDS Tissue Consortium brain gene array: two types of HIV-associated neurocognitive impairment. PLoS One 2012;7:e46178

22. Zink MC, Suryanarayana K, Mankowski JL, et al. High viral load in the cerebrospinal fluid and brain correlates with severity of simian immunodeficiency virus encephalitis. J Virol 1999;73:10480–8

23. Schefe JH, Lehmann KE, Buschmann IR, et al. Quantitative real-time RT-PCR data analysis: current concepts and the novel “gene expression’s CT difference” formula. J Mol Med (Berl) 2006;84:901–10

24. Meulendyke KA, Ubaida-Mohien C, Drewes JL, et al. Elevated brain monoamine oxidase activity in SIV- and HIV-associated neurological disease. J Infect Dis 2014;210:904–12

25. Knight AC, Brill SA, Queen SE, et al. Increased Microglial CSF1R Expression in the SIV/Macaque Model of HIV CNS Disease. J Neuropathol Exp Neurol 2018;77:199–206

26. Stein-O’Brien GL, Clark BS, Sherman T, et al. Decomposing Cell Identity for Transfer Learning across Cellular Measurements, Platforms, Tissues, and Species. Cell Syst 2019;8:395–411 e8

27. Saha RN, Pahan K. Differential regulation of Mn-superoxide dismutase in neurons and astroglia by HIV-1 gp120: Implications for HIV-associated dementia. Free Radic Biol Med 2007;42:1866–78

28. Sugawara T, Noshita N, Lewen A, et al. Overexpression of copper/zinc superoxide dismutase in transgenic rats protects vulnerable neurons against ischemic damage by blocking the mitochondrial pathway of caspase activation. J Neurosci 2002;22:209–17

29. Hendrickx DAE, van Eden CG, Schuurman KG, et al. Staining of HLA-DR, Iba1 and CD68 in human microglia reveals partially overlapping expression depending on cellular morphology and pathology. J Neuroimmunol 2017;309:12–22

30. Pandey HS, Seth P. Friends Turn Foe-Astrocytes Contribute to Neuronal Damage in NeuroAIDS. J Mol Neurosci 2019;69:286–97

31. Raghavan R, Stephens EB, Joag SV, et al. Neuropathogenesis of chimeric simian/human immunodeficiency virus infection in pig-tailed and rhesus macaques. Brain Pathol 1997;7:851–61

32. Mankowski JL, Queen SE, Tarwater PJ, et al. Elevated peripheral benzodiazepine receptor expression in simian immunodeficiency virus encephalitis. J Neurovirol 2003;9:94–100

33. Petito CK, Cho ES, Lemann W, et al. Neuropathology of acquired immunodeficiency syndrome (AIDS): an autopsy review. J Neuropathol Exp Neurol 1986;45:635–46

34. Brack-Werner R. Astrocytes: HIV cellular reservoirs and important participants in neuropathogenesis. AIDS 1999;13:1–22

35. Sabri F, Tresoldi E, Di Stefano M, et al. Nonproductive human immunodeficiency virus type 1 infection of human fetal astrocytes: independence from CD4 and major chemokine receptors. Virology 1999;264:370–84

36. McCarthy M, He J, Wood C. HIV-1 strain-associated variability in infection of primary neuroglia. J Neurovirol 1998;4:80–9

37. Di Rienzo AM, Aloisi F, Santarcangelo AC, et al. Virological and molecular parameters of HIV-1 infection of human embryonic astrocytes. Arch Virol 1998;143:1599–615

38. Churchill MJ, Wesselingh SL, Cowley D, et al. Extensive astrocyte infection is prominent in human immunodeficiency virus-associated dementia. Ann Neurol 2009;66:253–8

39. Ko A, Kang G, Hattler JB, et al. Macrophages but not Astrocytes Harbor HIV DNA in the Brains of HIV-1-Infected Aviremic Individuals on Suppressive Antiretroviral Therapy. J Neuroimmune Pharmacol 2019;14:110–9

40. Beck SE, Queen SE, Metcalf Pate KA, et al. An SIV/macaque model targeted to study HIV-associated neurocognitive disorders. J Neurovirol 2018;24:204–12

41. Zelko IN, Mariani TJ, Folz RJ. Superoxide dismutase multigene family: a comparison of the CuZn-SOD (SOD1), Mn-SOD (SOD2), and EC-SOD (SOD3) gene structures, evolution, and expression. Free Radic Biol Med 2002;33:337–49

42. Ishihara Y, Takemoto T, Itoh K, et al. Dual role of superoxide dismutase 2 induced in activated microglia: oxidative stress tolerance and convergence of inflammatory responses. J Biol Chem 2015;290:22805–17

43. Bagashev A, Sawaya BE. Roles and functions of HIV-1 Tat protein in the CNS: an overview. Virol J 2013;10:358

44. Budka H, Costanzi G, Cristina S, et al. Brain pathology induced by infection with the human immunodeficiency virus (HIV). A histological, immunocytochemical, and electron microscopical study of 100 autopsy cases. Acta Neuropathol 1987;75:185–98

45. Watkins BA, Dorn HH, Kelly WB, et al. Specific tropism of HIV-1 for microglial cells in primary human brain cultures. Science 1990;249:549–53

46. Koenig S, Gendelman HE, Orenstein JM, et al. Detection of AIDS virus in macrophages in brain tissue from AIDS patients with encephalopathy. Science 1986;233:1089–93

47. Boven LA, Gomes L, Hery C, et al. Increased peroxynitrite activity in AIDS dementia complex: implications for the neuropathogenesis of HIV-1 infection. J Immunol 1999;162:4319–27

48. Ances BM, Roc AC, Korczykowski M, et al. Combination antiretroviral therapy modulates the blood oxygen level-dependent amplitude in human immunodeficiency virus-seropositive patients. J Neurovirol 2008;14:418–24

49. Akay C, Cooper M, Odeleye A, et al. Antiretroviral drugs induce oxidative stress and neuronal damage in the central nervous system. J Neurovirol 2014;20:39–53

50. Robertson K, Liner J, Meeker RB. Antiretroviral neurotoxicity. J Neurovirol 2012;18:388–99

51. Elzi L, Erb S, Furrer H, et al. Adverse events of raltegravir and dolutegravir. AIDS 2017;31:1853–8

52. Montenegro-Burke JR, Woldstad CJ, Fang M, et al. Nanoformulated Antiretroviral Therapy Attenuates Brain Metabolic Oxidative Stress. Mol Neurobiol 2019;56:2896–907

