## Supplementary figures and images for "UPREGULATION OF SUPEROXIDE DISMUTASE 2 BY ASTROCYTES IN THE SIV/MACAQUE MODEL OF HIV-ASSOCIATED NEUROLOGIC DISEASE"

### Supplemental Figure 1

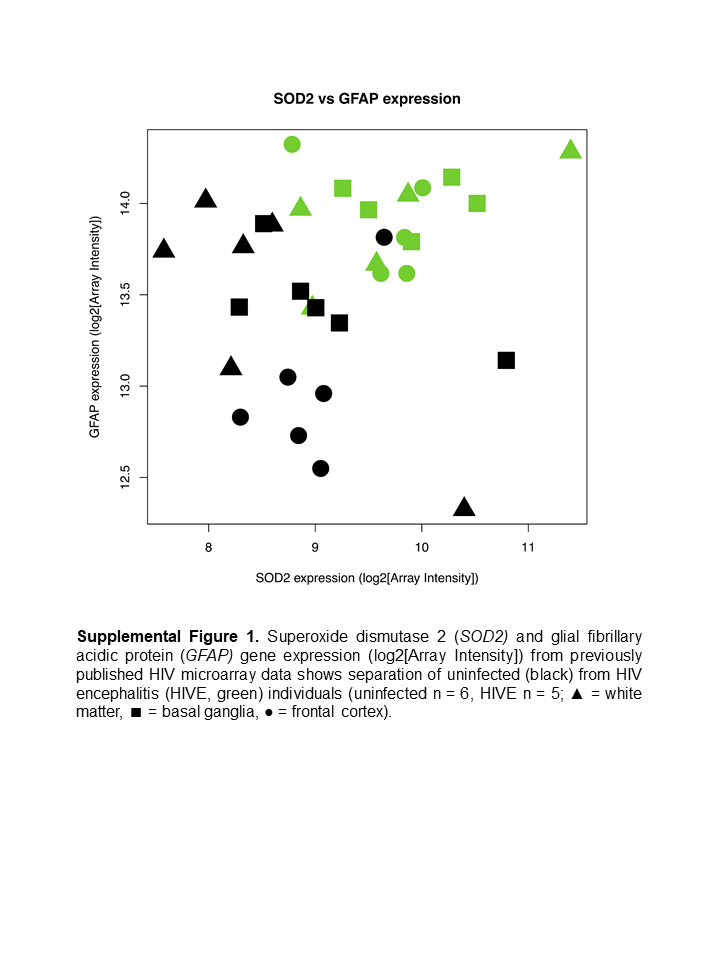
