## Supplemental Table 1 for "UPREGULATION OF SUPEROXIDE DISMUTASE 2 BY ASTROCYTES IN THE SIV/MACAQUE MODEL OF HIV-ASSOCIATED NEUROLOGIC DISEASE"

| Gene | SIV *p* value | HIV *p* value | PC1 weight |
| --- | --- | --- | --- |
| B2M | 4.57E-05 | 2.31E-08 | 0.735 |
| GFAP | 5.48E-04 | 2.31E-04 | 0.543 |
| SOD2 | 4.57E-05 | 1.35E-03 | 0.297 |
| UBC | 4.57E-05 | 2.59E-01 | 0.157 |
| MX1 | 4.46E-04 | 1.87E-07 | 0.122 |
| SLC1A2 | 6.22E-03 | 7.89E-01 | -0.095 |
| SIV17E | 4.22E-04 | NA | 0.082 |
| SNAP25 | 2.66E-02 | 2.15E-01 | -0.063 |
| MBP | 2.37E-01 | 2.15E-01 | -0.058 |
| TSPAN7 | 6.22E-03 | 6.30E-01 | -0.055 |
| CXCL10 | 4.57E-05 | 4.42E-01 | 0.041 |
| YWHAZ | 1.37E-03 | 7.35E-01 | -0.040 |
| SLC1A3 | 5.45E-02 | 3.64E-02 | 0.036 |
| STAT1 | 4.57E-05 | 9.80E-07 | 0.031 |
| GLUL | 4.39E-03 | 5.19E-03 | 0.027 |
| APP | 8.68E-04 | 5.32E-01 | -0.027 |
| GAPDH | 8.55E-03 | 3.25E-01 | -0.027 |
| ACTB  DLG4  CAMK2A | 2.66E-02  5.45E-02  1.01E-01 | 7.08E-01  8.73E-01  7.08E-01 | 0.019  -0.017  -0.017 |

**Supplemental Table 1**. Top 20 genes with the largest Principal Component 1 (PC1) absolute value. Reported *p* values are Mann-Whitney tests showing differential expression of the genes from uninfected macaques versus SIV-infected macaques with encephalitis (SIV *p* value), and uninfected individuals versus HIV-infected individuals with encephalitis (HIV *p* value). Negative PC1 weights indicate a downregulation of the gene in individuals with encephalitis, while positive weights indicate an upregulation of the gene in those individuals.
